# Single-lipid dynamics in phase-separated supported lipid bilayers

**DOI:** 10.1101/2020.05.28.121830

**Authors:** Xinxin Woodward, Christopher V. Kelly

## Abstract

Phase separation is a fundamental organizing mechanism on cellular membranes. Lipid phases have complex dependencies on the membrane composition, curvature, tension, and temperature. Single-molecule diffusion measures a key characteristic of membrane behavior and relates to the effective membrane viscosity. Lipid diffusion rates vary by up to ten-fold between liquid-disordered (*L_d_*) and liquid-ordered (*L_o_*) phases depending on the membrane composition, measurement technique, and the surrounding environment. This manuscript reports the lipid diffusion on phase-separated supported lipid bilayers (SLBs) with varying temperature, composition, and lipid phase. Lipid diffusion is measured by single-particle tracking (SPT) and fluorescence correlation spectroscopy (FCS) via custom data acquisition and analysis protocols that apply to diverse membranes systems. We demonstrate agreement between FCS and SPT analyses with both the single-step length distribution and the mean squared displacement of lipids with significant immobile diffusers. Traditionally, SPT is sensitive to diffuser aggregation, whereas FCS largely excludes aggregates from the reported data. Protocols are reported for identifying and culling the aggregates prior to calculating diffusion rates via SPT. With aggregate culling, all diffusion measurement methods provide consistent results. With varying membrane composition and temperature, we demonstrate the importance of the tie-line length that separates the coexisting lipid phases in predicting the differences in diffusion between the *L_d_* and *L_o_* phases.

**HIGHLIGHTS:**

- Lipid diffusion varies with the lipid phases, temperature, and aggregation
- Aggregate culling yields consistent measurements from single-particle tracking and fluorescence correlation spectroscopy
- Membrane with higher cholesterol content or at low temperature have more aggregates
- A more variation in the diffusion rates occurred between the coexisting lipid phases at low temperatures and low cholesterol content

## 1) INTRODUCTION

Cell plasma membranes are often modeled as a two dimensional fluid with lipid phase separation (Pike, 2006; Pralle et al., 2000; Simons and Ikonen, 1997). Lipid phases are hypothesized to be critical for cell functions such as protein sorting, cell signaling, and membrane budding (Fessler and Parks, 2011; Hurley et al., 2010; Simons and Toomre, 2000). Model membranes enable detailed analyses of coexisting liquid phases by connecting the membrane composition with biophysical observables, such as viscosity, bilayer thickness, and fluctuation analyses (Kiessling et al., 2015; Veatch and Keller, 2005). However, anomalous diffusion, lipid confinement, and nanodomains complicate the measurement of the membrane properties while revealing heterogeneity in lipid behavior.

Model membranes can phase separate into a liquid-ordered phase (*L_o_*) and a liquid-disordered phase (*L_d_*) when composed of a mixture of three lipid types: a phospholipid with a high melting temperature, a phospholipid with low melting temperature, and a sterol (Veatch and Keller, 2002). Lipids that have a high melting temperature tend to have longer and more saturated acyl tails while concentrating in the *L_o_* phase. Lipids that have a low melting temperature commonly have shorter, unsaturated tails while concentrating in the *L_d_* phase. The most commonly used sterol in model membranes is cholesterol and it slightly sorts to the *L_o_* phase. Separate condensed complexes that are composed of cholesterol and DPPC may further affect the lipid dynamics, but do not appear with diffraction-limited miroscopy (McConnell and Radhakrishnan, 2003; Radhakrishnan and McConnell, 2005).

Common model membrane compositions and temperatures yield the *L_o_* phase being up to 0.8 nm ticker (Bleecker et al., 2016; Chiantia et al., 2006b, 2006a; Lin et al., 2007), having up to 3x greater bending rigidity (Baumgart et al., 2003; Dimova, 2014; Gracià et al., 2010; Kollmitzer et al., 2015), and providing up to 10x greater effective viscosity (Kahya et al., 2003; Scherfeld et al., 2003) than the *L_d_* phase. Increasing the sample temperature shortens the tie-lines and causes the *L_o_* and *L_d_* phases to become more similar in composition and behavior (Fig. 1). If the temperature is above the miscibility transition temperature (*T_m_*), a single liquid phase (*L*) exists. The phases reemerge when the membrane temperature drops below *T_m_* with an interplay of phase nucleation and growth dynamics. With increasing temperature, the diffusion rate of the lipids increases (Almeida et al., 1992; Bag et al., 2014; Jacobson et al., 1981; Tamm, 1988; Wu et al., 1977). If the temperature of a phase-separated membrane is increased above *T_m_*, then the two populations of lipids with over a 3x differnces in diffusion rate may become a single population (Filippov et al., 2004; Lindblom et al., 2006).

**Figure 1:**
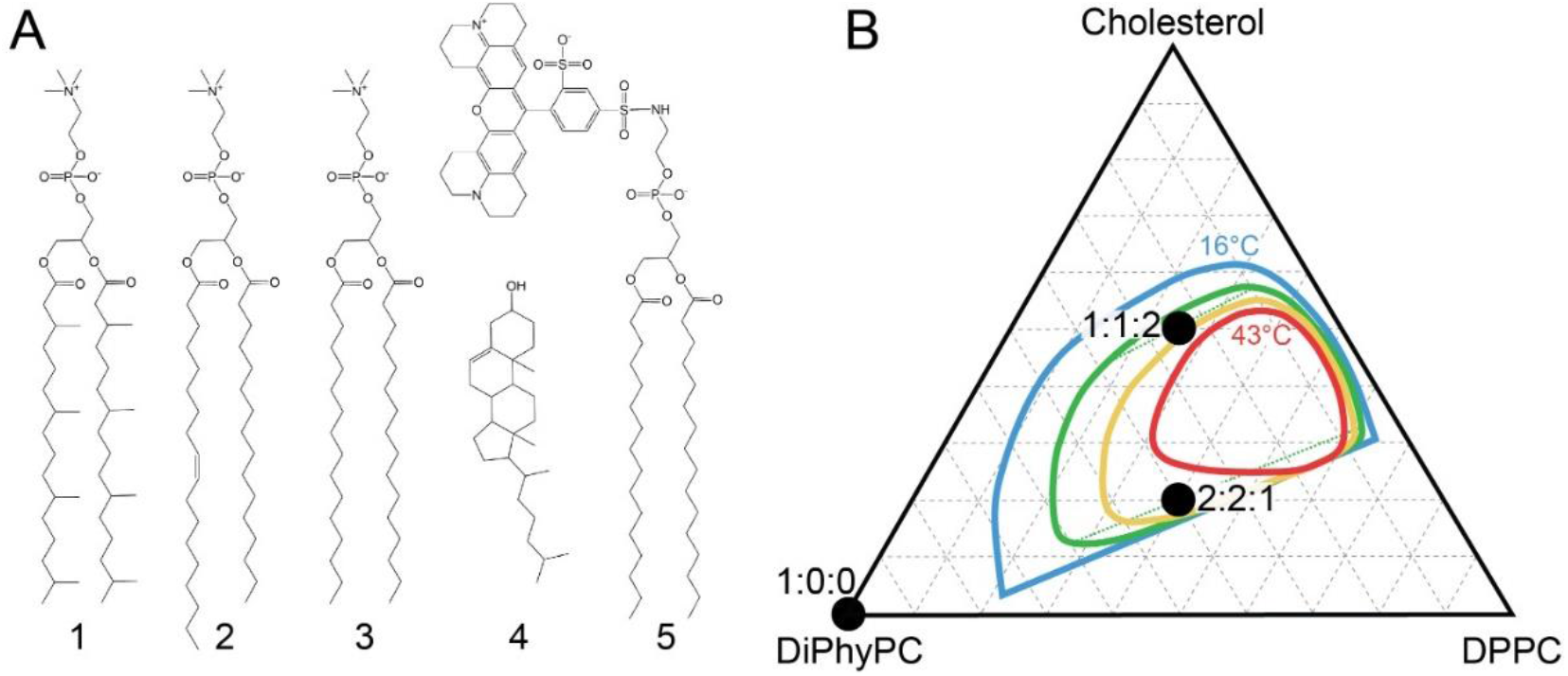
(A) The lipids used in this study include DiPhyPC (1), POPC (2), DPPC (3), cholesterol (4), and DPPE-TR (5). (B) Mixtures of DiPhyPC, DPPC, and cholesterol display two coexisting liquid phases within the indicated regions that vary with temperature. The DiPhyPC:DPPC:cholesterol molar ratios used in this study are shown (*black dots*). The 16°C (*blue*) and 43°C (*red*) phase boundaries were measured previously via fluorescence microscopy of GUVs (Veatch et al., 2006). The approximate boundary of liquid phase coexistence at 25°C (*green*) and 34°C (*yellow*) for SLBs are also shown. The approximate tie-line at 25°C (*green dotted*) is longer for the composition of 2:2:1 than 1:1:2, indicating a greater difference between the liquid phases for 2:2:1 vs. 1:1:2.

Varying membrane preparations and analysis methods provide differing resolution to heterogenious populations and non-Brownian diffusion. Lipids may form clusters and demonstrate confined motion due to interactions with the supporting substrates (Beckers et al., 2020; Hsieh et al., 2014; Spillane et al., 2014; Wawrezinieck et al., 2005), nonspecific lipid-lipid interactions (Jan Akhunzada et al., 2019; Scherer et al., 2015; Spillane et al., 2014), specific condensed complexes (McConnell and Radhakrishnan, 2003; Radhakrishnan and McConnell, 2005), lipid crosslinking (Štefl et al., 2012), incomplete bilayer formation (Coker et al., 2019), and phase-associated nanodomains (Sodt et al., 2014; Wu et al., 2016). For example, even bilayers with ≤2 mol% GM1 form isolated nanodomains within larger lipid phases (Sun et al., 2015; Yuan et al., 2002; Yuan and Johnston, 2001). Robust methods to characterize lipid clustering or aggregation are needed to resolve the varying modes of diffusion and influences of lipid behavior.

Two complementary techniques to study single-molecule diffusion are fluorescence correlation spectroscopy (FCS) and single-particle tracking (SPT) (Harwardt et al., 2018). FCS uses less total light power compared to SPT (>10 mW vs. <10 μW, respectively), and FCS measures higher fluorescent lipid concentrations than SPT (>200 lipids/μm^2^ vs. <6 lipids/μm^2^, respectively). However, FCS is inherently blind to immobile diffusers and rare events are difficult to detect (Guo et al., 2012). FCS has a spatial resolution conventionally limited by the diffraction-limited beam waist size (*i.e*., 250 nm). The spatial resolution of SPT depends on the single-molecule localization precision (*i.e*., 1-20 nm), the step length between imaging frames (*i.e*., 10-300 nm), and the analysis method. SPT may provide a sub-diffraction-limited mapping of the spatial variations in the effective viscosity, nano-domains, and aggregates. For example, FCS shows little effects from nanoscale membrane curvature relative to the curvature-dependent lipid mobility change in simulations (Kabbani et al., 2017). By mapping the single-lipid step lengths to locations on the membrane, the lipid mobility relative to membrane curvature has been analyzed (Kabbani and Kelly, 2017a; Woodward et al., 2018; Woodward and Kelly, 2020).

This manuscript details complementary analysis procedures for SPT trajectories to map the lipid diffusion rate across a sample while lipid aggregation is present. Aggregate identification and data culling were performed to study single-lipid diffusion with varying the temperature, lipid composition, and coexisting lipid phases. Consistency was demonstrated between FCS and SPT, including analyzing the SPT results with MSD or single-step-length distributions. In all conditions, faster diffusion was observed for higher temperatures, and greater differences between coexisting phases were observed when the tie-lines were longer. The protocols developed here are directly applicable to mapping lateral heterogeneities in effective membrane viscosities, such as those created by nanoscale curvature (Woodward and Kelly, 2020).

## 2) MATERIALS AND METHODS

### 2.1) GUV formation

Quasi-one component bilayers composed of diphytanoylphosphatidylcholine (DiPhyPC; Avanti Polar Lipids) or palmitoyloleoylphosphatidylcholine (POPC, Avanti Polar Lipids) were used to examine the liquid phase of a one component membrane (*L_α_*) at room temperature (Fig. 1A). DiPhyPC was used more frequently than POPC in this manuscript because DiPhyPC provides saturated, branched acyl tails provide both resistance to oxidization and a low melting transition temperature <-120°C (Lindsey et al., 1979). DiPhyPC was combined with dipalmitoylphosphatidylcholine (DPPC; Avanti Polar Lipids) and cholesterol (Avanti Polar Lipids) to study phase-separated SLBs. A fluorescent lipid, dipalmitoylphosphoethanolamine-Texas Red (DPPE-TR, Life Technologies), was used to label the *L_d_* domain with a total concentration of 0.1 mol% in all membranes. We used three compositions with 1:0:0, 2:2:1, and 1:1:2 molar ratios of DiPhyPC:DPPC:cholesterol. The buffers were created from Milli-Q water with a resistivity of 18 mΩ. All other chemicals were bought from Sigma Aldrich and used without further purification.

All samples were created by the fusion of giant unilaminar vesicles (GUVs) on a glass coverslip. Our GUV making protocol was adapted from previous reports (Veatch, 2007). Lipids were combined at the desired ratio in chloroform with a concentration of 5 mg/mL and dried onto electrically conducting indium tin oxide coated glass plates. A trimmed silicon sheet was added between the plates and stabilized by clips to create an incubation chamber. The chamber was filled with 200 mM sucrose and exposed to an AC voltage with V_rms_ of 3 V at 10 Hz for 1 hr at 55°C. The resulting GUV solution had a concentration of 13 mg lipids per mL and was stored at 55°C until use. The GUVs were used within 2 days of their electroformation.

### 2.2) Substrate preparation

All SLBs were formed and imaged on 35 mm diameter glass-bottom dishes (MatTek). Dishes were initially rinsed with ethanol, dried by a nitrogen gas stream, and placed in air plasma (Harrick Plasma) for 10 s to create a hydrophilic surface. 20 μL of 50 mM CaCl_2_ was deposited on the dish and dried on a hot plate at 35°C for 10 min. At least three SLB patches from each of two GUV batches were used for each condition.

### 2.3) Supported-lipid bilayer formation

GUVs were cooled to 4°C prior to SLB formation. 5 μL of 4°C GUVs and 50 μL of 4°C Milli-Q water were sequentially deposited on the room temperature glass-bottom dish. The dishes were maintained at 4°C for 15 min before rinsed gently with 5 mL of 4°C 200 mM sucrose. We note that the dishes were at room temperature prior to the addition of the GUV solution. If the sample dish was cooled to 4°C before GUV deposition, the condensation on glass reduced GUV fusion. The resulting SLB patches had lipid phase domains with similar size and shape as seen on the GUVs prior to fusion (Fig. 2).

**Figure 2:**
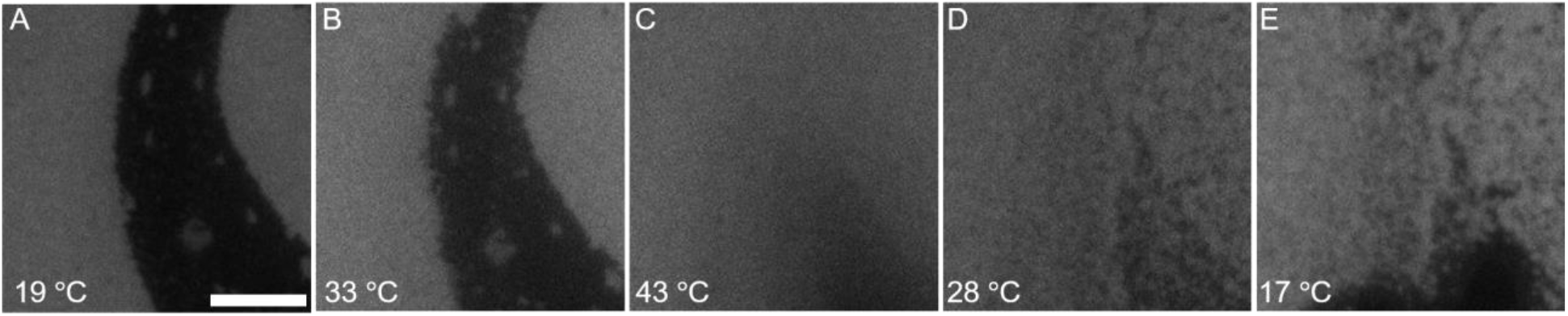
The coexisting phases in an SLB with a 1:1:2 molar ratio of DiPhyPC:DPPC:cholesterol mix upon heating and reform upon cooling. Below 35°C, the bilayer is phase-separated into *L_d_* (*bright*) and *L_o_* (*dim*) phases due to differential partitioning of the fluorescent DPPE-TR. These panels are shown in the order in which they were acquired with 50 min between frames. Scale bar, 5 μm.

### 2.4) Temperature control

A Peltier temperature control dish holder (QE-1HC, Warner Instruments) was used with custom, insulated dish cover, LabVIEW control, and a thermocouple holder. The thermocouple was positioned <0.5 mm above the center of the glass coverslip for real-time temperature feedback. The temperature was changed by 0.5°C/min and remained at each set temperature for ≥30 min before data aquisiton. The dish holder was initially set to 10°C. When the temperature was set to 10°C, 30°C, or 45°C, the resulting measured temperature was 17 ± 3°C, 27 ± 1°C, and 37 ± 1°C, respectively. The dishes were never heated above 45°C due to limitations in our optical system.

### 2.5) Single-fluorophore imaging

The optical setup included an inverted IX83 microscope with a 100x, 1.49 NA objective (Olympus), a 2x emission path magnification (OptoSplit, Cairn Research), and an iXon 897-Ultra EMCCD camera (Andor Technology). A Hg lamp with an excitation filter (BrightLine single-band, Semrock) provided wide-field fluorescence illumination for the diffraction-limited images. CUBE diode lasers with wavelengths of 405 and 488 nm (Coherent) and a 561 nm Sapphire laser (Coherent) were used for single-fluorophore excitation. The laser light passed through a clean-up filter (zet405/488/561/647x, Chroma Technology), reflected from a quad-band dichroic mirror (zt405/488/561/647rpc, Chroma Technology), and transmitted into the objective to excite the sample. The fluorescence emission was isolated via emission filters (BrightLine single-band filter, Semrock) and a 4-band notch laser filter (zet405/488/561/640 m, Chroma Technology). SOLIS software (Andor Technology) was used to acquire images and movies with a 128 pixels x 128 pixels region of interest in kinetic model and an EM gain of 150. Videos of single-fluorophores binking were acquired at 537 Hz with ≥20,000 frames per sample.

### 2.6) Single-fluorophore localization

The videos of optically isolated fluorescent lipids were analyze with Fiji plug-in ThunderSTORM (Ovesný et al., 2014; Schindelin et al., 2012). Via ThunderSTORM, each bright spot in the movies was fit with a 2D Gaussian function to identify the location, intensity, fit width, localization uncertainty (*σ_r_*), and brightness of each fluorophore. Only the fluorophore localizations with intensity >100 photons, Gaussian fit width >15 nm, and *σ_r_* <45 nm were kept for further analysis. ThunderSTORM reported *σ_r_* = 24 ± 1 nm for the remaining localizations.

### 2.7) Aggregate removing method

Aggregated localizations were identified via spatial autocorrelation analysis. The localizations were two-dimensionally histogrammed to create a reconstructed image, 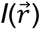. Spatial autocorrelation functions, *g_s_*(*r*), were calculated with fast Fourier transforms (FFTs) according to

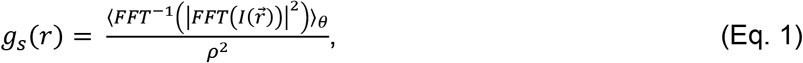

for which *ρ* is the average localization density, and averaging was performed over the azimuthal angle (*θ*), as done previously (Veatch et al., 2012). We calculated *g_s_* from the experimental and simulated localization data to yield *g_exp_* and *g_sim_*, respectively. The simulated localizations were randomly distributed over the same membrane shape that was observed experimentally to provide an aggregate-free normalization.

The average aggregate size and the number of localizations per aggregate (*N*) were calculated by identifying the exponential decay length scale and Eq. 2, respectively, as done previously (Shelby et al., 2013).

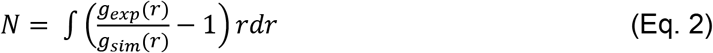

Aggregate removal included the grouping of all single-fluorophore localizations to 3D voxels of time and *xy*-position. When the number of localizations per voxel was above a density threshold (*ρ_th_*), the localizations were assessed to be part of an aggregate and removed from subsequent analysis. *ρ_th_* was varied to remove between 0 to 80% of all localizations for testing. For each *ρ_th_*, the remaining experimental localizations were used to calculate *N*. Decreasing *ρ_th_* resulted in decreasing *N*, as expected. The *ρ_th_* value that was chosen for aggregate-removed diffusion analysis was the maximum *ρ_th_* value that yielded *N* ≤ 3.

### 2.8) Single-molecule trajectory analysis

The remaining localizations after aggregate removal were linked through u-track (Jaqaman et al., 2008) with a maximum linking radius of 400 nm. Trajectories with more steps yielded a slower diffusion compared to shorter trajectories, as observed previously (Saxton, 1997). The longer trajectories were possibly from oligomers, as would be expected to have slower diffusion than single lipids. Therefore, trajectories longer than 32 steps were removed from all diffusion studies whenever aggregates were removed. These aggregate and long-trajectory removal methods resulted in the average trajectory consisting of 6 ± 4 steps.

#### 2.8.1) Rayleigh distribution analysis

The single-fluorophore step lengths (*v*) were fit to Rayleigh distribution (*R*) to find the diffusion coefficient (*D_fit_*), as done previously (Cheney et al., 2017; Kabbani et al., 2017; Kabbani and Kelly, 2017b; Knight et al., 2010; Woodward et al., 2018),

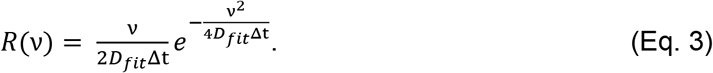

*D_fit_* was corrected for imaging blur and localization uncertainty to yield the diffusion coefficient from this single-step length analysis (*D_RD_*) (Berglund, 2010; Lagerholm et al., 2017; Qian et al., 1991).

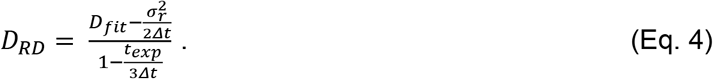

The difference between the *D_fit_* and *D_RD_* values depends on the single-frame exposure time (*t_exp_*), the time between frames (*Δt*), and localization uncertainty (*σ_r_*) exported by ThunderSTORM.

#### 2.8.2) Mean squared displacement analysis

The mean-squared displacement (MSD) as a function of *Δt* was calculated for all samples (Wu et al., 1977). *D_MSD_* was found by fitting the first two time points (*i.e., Δt* = 1.9 and 3.8 ms) of the localization uncertainty-corrected MSD vs. *Δt*,

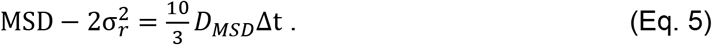

By using *σ_r_* exported from ThunderSTORM rather than incorporating it as an unknown in the fitting routine, *D_MSD_* and *D_RD_* could be directly compared without using the y-intercept of the MSD vs. *Δt* fit before performing the Rayleigh distribution analysis.

## 2.9) Fluorescence correlation spectroscopy

Custom fluorescence correlation spectroscopy (FCS) was incorporated into an inverted IX71 microscope with a 40x, 1.3 NA objective (Olympus). The excitation laser was a super-continuum fiber laser (SC-Pro, YSL photonics). The long-wavelength component (>650 nm) was removed before the optical excitation were chromatically filtered (BrightLine FF01-561/14-25, Semrock), expanded, and focused by the microscope objective onto the sample. The laser power entering the microscope objective was 1.6 μW. A beam waist of 180 ± 20 nm was formed on the SLB. The fluorescence emission was filtered (ZET442/514/561m, Chroma) and collected on an sCMOS camera (Zyla, Andor Technology). Each camera frame was acquired for 1 ms at 900 Hz frame rate during a 10-sec acquisition. The mean intensity of each camera frame was calculated to be the intenisity (*I*) of the emission. The temporal auto-correlation function (*g_FCS_*) was calculated from intensity vs. time according to

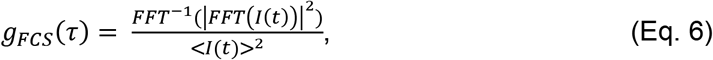

where <> represents the time average. All FCS experiments were performed at room temperature (25 ± 2°C). Z-scan FCS was performed to optimize the focus and to ensure the minimum beam waist was consistently obtained. Five locations from each of two sample dishes were measured for each condition.

## 3) RESULTS

### 3.1) Creating phase-separated SLBs

We created phase-separated SLBs with 1:1:2 and 2:2:1 molar ratios of DiPhyPC:DPPC:cholesterol. The phases were identified by the differential partitioning of the fluorescent lipid, DPPE-TR, such that higher concentrations of DPPE-TR were found in more disordered phases. All phases appeared to be liquid with rapid diffusion and merging of the phases in GUVs. The phases on the SLBs were similar in size and shape to those on the GUVs but immobile on the time scale on our experiments. The varying membrane compositions between GUVs in a single preparation resulted in varying *T_m_* and DPPE-TR partition coefficients for separate SLB patches in a single sample dish. The 1:1:2 SLBs had higher cholesterol content and a lower *T_m_* than the 2:2:1 SLBs (Fig. 1B).

During the optimization of the SLB formation method, the importance of deposition temperature, buffer, and substrate preparation were noted. Our initial experiments yielded SLBs that contained membrane defects (*i.e*., holes in the bilayer or abundant fused nanoscale vesicles) or did not maintain the large-scale phase separation that was seen on the GUVs. The keys parameters for high-quality SLB formation were to cool the GUVs and the buffers before GUV fusion in sucrose-rich solution with trace CaCl_2_, as detailed in the Methods section. For example, when we direction transferd GUVs from the 55°C incubator to the room-temperature glass coverslip, the SLBs consistently demonstrated a phase separation with the L_d_ phase at the center of the SLB patch and L_o_ phase at the perimeter as the coverslip. SLB formation from GUV fusion in the presence of sucrose solution resulted in smoother SLBs than when an ionic buffer was used; vesicles of sub-micron diameter covered the SLB patches when phosphate buffered saline was present during SLB formation.

### 3.2) Temperature-induced phase changes

A convenient method for dynamically varying the differences between the phases in a single sample is to change the sample temperature. As the temperature is increased, the fluorescence emission from the *L_d_* and *L_o_* phases becomes more similar (Fig. 2), as seen previously (Gunderson and Honerkamp-Smith, 2018). This is consistent with an increasing temperature resulting in less compositional difference and a shorter tie-line separating the coexisting phases. When the temperature is held slightly above *T_m_*, the two phases mix into a single liquid phase (*L*) with composition and material properties between the prior *L_d_* and *L_o_* phases.

The majority of SLBs with a 1:1:2 molar ratio of DiPhyPC:DPPC:cholesterol demonstrated phase mixing with a uniform fluorescence emission across the sample after the temperature was maintained at 40°C for 30 min. However, the large domains of area >100 μm^2^ were unable to mix fully during this time and maintained partial phase separation throughout our observation (Fig. 2C).

As the temperature was decreased from 40°C to 30°C for 1:1:2 SLBs, domains became optically identifiable and enlarged within 5 min. However, the domains did not grow larger than nanoscale in diameter during our 30 min of observation at each temperature (Fig. 2D). The locations of new domain formation upon cooling were typically random and not dependent on the prior domain locations. However, large domains that were not fully mixed at 40°C demonstrated *L_o_* phase domains preferentially forming at the prior *L_o_* regions.

SLBs with composition 2:2:1 molar ratio of DiPhyPC:DPPC:cholesterol had *T_m_* above 40°C. The coexisting *L_o_* and *L_d_* phases remained sharply defined after 120 min at 40°C. These cholesterol-poor membranes showed consistently large differences between the phases. The phase boundaries were always clearly defined, which is consistent with a large tie-line separating the *L_o_* and *L_d_* phases.

### 3.3) Aggregate identification and removal

The single-molecule localization data was reconstructed as super-resolution images with Voronoi diagrams. Voronoi diagrams are created by dividing the imaged area into polygons such that each polygon represents the area of the image that is closest to a particular single-molecule localization. The polygon area represents the localization density without binning biases and is colored appropriately. The lateral variations across the Voronoi images of DPPE-TR localizations in POPC bilayers was consistent with random sampling density variations expected from the total number of localizations acquired (Fig. 3a). However, non-random distributions of DPPE-TR localizations were observed in all DiPhyPC-containing bilayers (Fig. 3b). Unlike the super-resolution images of the POPC bilayer, the super-resolution images with DiPhyPC showed <70 nm diameter aggregates of dense localizations.

**Figure 3:**
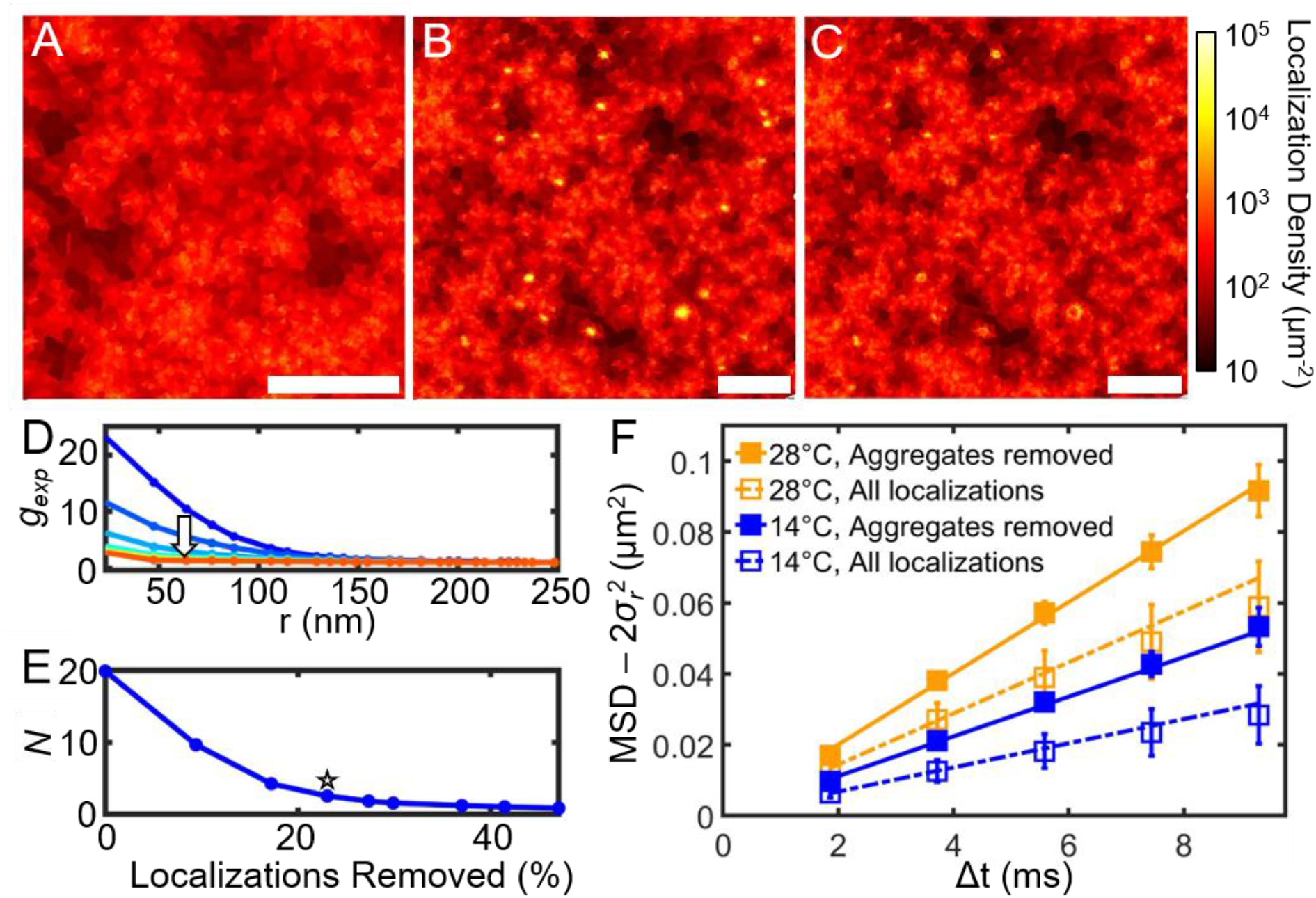
Aggregation was observed in SLBs with DiPhyPC but not POPC. A Voronoi diagram of DPPE-TR localizations in a POPC bilayer at 25°C shows randomly distributed localizations (A). Voronoi diagrams of DPPE-TR localizations in DiPhyPC bilayers at 14°C (B) without and (C) with aggregates removed. (D) Spatial autocorrelations of localizations with increasing *p_th_* (*arrow*). (E) The localizations per aggregate (*N*) and the resulting fraction of localizations removed for increasing *p_th_*. The greatest *p_th_* value that yielded *N* ≤ 3 (*star*) was used for aggregate-free analysis. (F) MSD for DPPE-TR in DiPhyPC SLBs with and without aggregates included at 14°C and 28°C. Error bars represent the standard error from at least three repeated measurements. (A-C) Scale bars, 1 μm.

Quantification of the aggregates was performed by calculating the spatial autocorrelation (*g_s_*) (Eq. 1). Random distributions of localizations yield *g_s_* = 1 with no increased probability of finding a localization in the proximity of other localizations. The presence of aggregates, however, appears in *g_s_* because localizations are more likely to be found close together. *g_s_* reveals the size and density of the average aggregate. The area under *g_s_* measures the number of localizations in an aggregate compared to the sample average localization density. The aggregates within DiPhyPC-containing SLBs were 63 ± 13 nm diameter and contained *N* = 20 ± 1 localizations each for all compositions tested (Fig. 3D and E).

A threshold density of localizations (*ρ_th_*) was used to identify and remove aggregation. All localizations from the regions of the SLB that displayed localization rates above *ρ_th_* were culled to provide an aggregate-free assessment of the single-lipid diffusion. Smaller values of *ρ_th_* resulted in smaller values of *N* from the remaining localizations. *ρ_th_* was set to be the largest value that provided *N* ≤ 3 for further analysis. *ρ_th_* varied between 430 – 82000 localizations μm^−2^ s^−1^ depending on the density of the aggregates, the experiment duration, the laser power, and the fluorophore density. Additionally, any localizations that were linked to be a trajectory lasting more than 32 steps was determined to be an outlier and removed from further analysis. After the aggregates and long trajectories were removed, *D_MSD_* increased in DiPhyPC SLBs by (1.7 ± 0.6)x at 14°C and (1.4 ± 0.3)x at 28°C (Fig. 3F).

Aggregates appeared in all DiPhyPC-containing membranes (*i.e*., 1:0:0, 1:1:2, and 2:2:1) and with a greater abundance when the tie-line between coexisting phases was longer (Fig. 4). As the temperature was increased, the number of aggregates present on the membrane decreased. As the cholesterol concentration in the membrane was increased, the number of aggregates present on the membrane increased. There was no significant difference in aggregate percentage between *L_d_* and *L_o_* phases.

**Figure 4:**
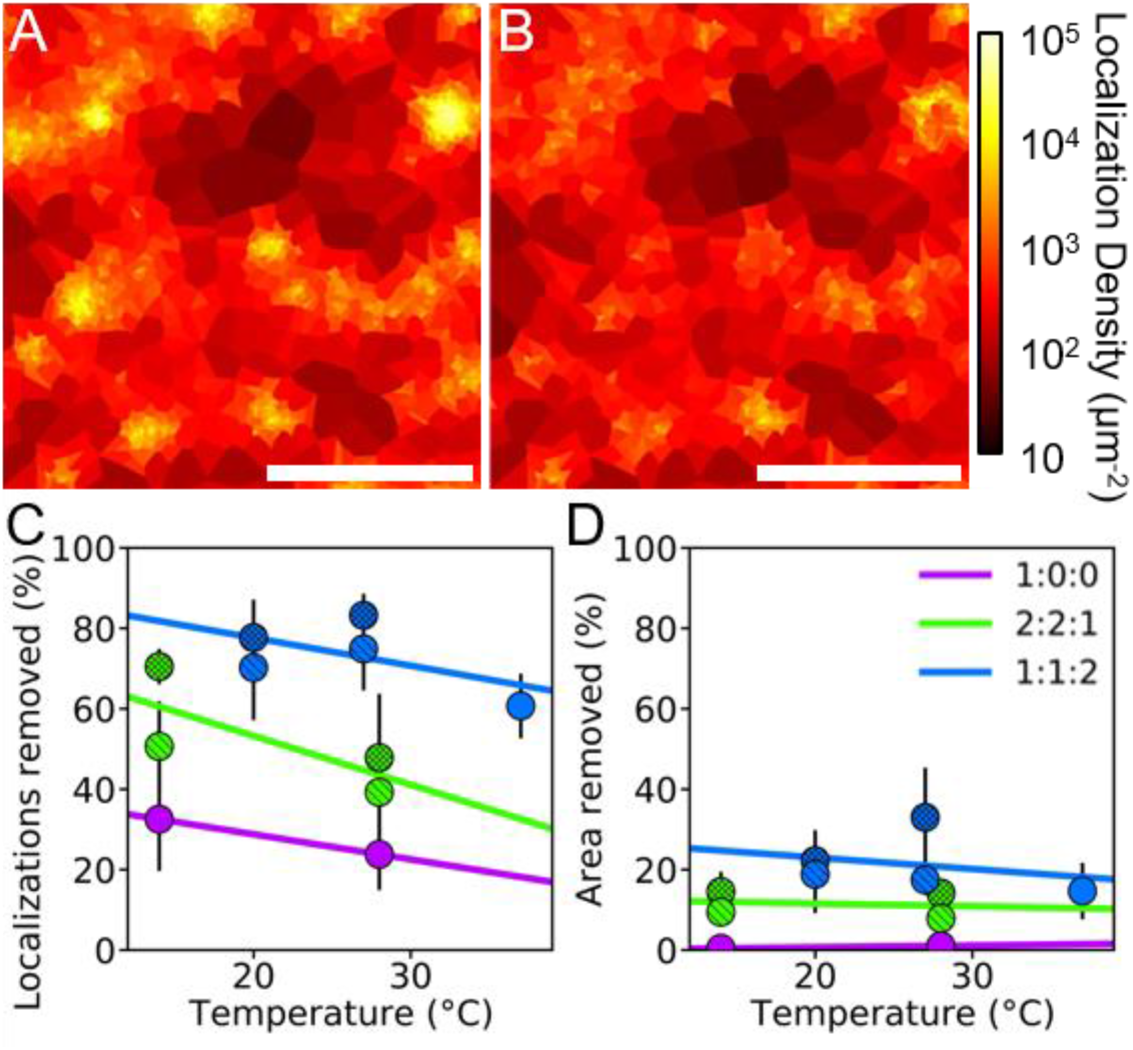
Aggregates were present in SLBs with DiPhyPC. Representative Voronoi diagrams of DPPE-TR localizations in a 1:1:2 SLB L_d_ at 20°C (A) without and (B) with aggregates removed. Scale bars, 0.5 μm. The fraction of the removed (C) localizations and (D) SLB area shows more aggregates removed when the tie-lines were longer. Marker hatching indicates the lipid phase: backslash = *L_o_*; crosshatching = *L_d_*; empty = *L* or *L_a_*. Linear fits are shown to guide the eye. Error bars represent the standard error of at least three repeated measurements.

### 3.4) Diffusion vs. phase

*D_MSD_* and *D_RD_* after aggregate removal were measured for all membrane compositions and temperatures (Table S1). The diffusion difference between the *L_d_* and *L_o_* phases was greater for membranes with less cholesterol in which the tie-lines were longer. For example, *D_MSD_* from the *L_d_* phase was (1.8 ± 0.6)x greater than that of the *L_o_* phase at both 14°C and 28°C for 2:2:1 SLBs (Fig. 5A). The higher cholesterol, 1:1:2 SLBs displayed no significant difference in *D_MSD_* from the *L_d_* and *L_o_* phases (Fig. 5B).

**Figure 5:**
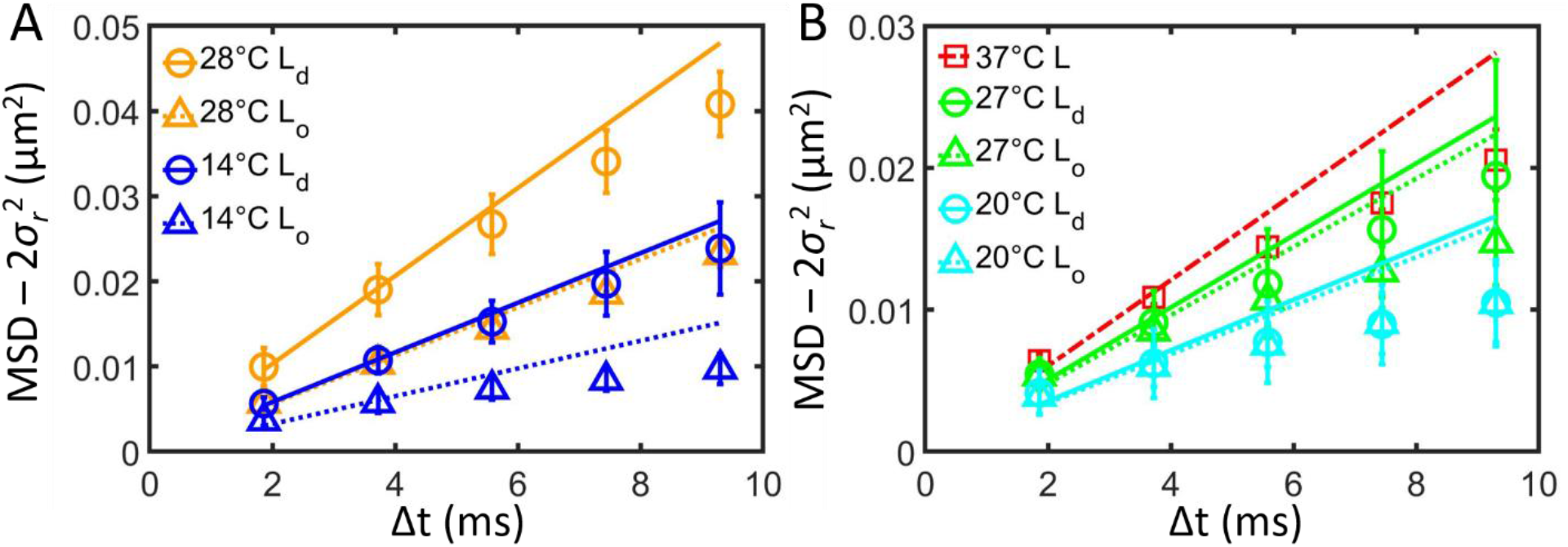
MSD vs. Δt for (A) 2:2:1 and (B) 1:1:2 SLBs demonstrate the importance of temperature and phase in single-lipid diffusion. The measured MSD values were corrected by the localization uncertainty reported by ThunderSTORM (*σ_r_*) such that the resulting linear fits have a y-intercept of zero. Fits were performed for the short time steps (Δt = 1.9-3.8 ms). Error bars represent the standard error of the means with at least three repeats.

### 3.5) Single-particle tracking vs. FCS

FCS provides a measure of single-lipid diffusion averaged over a diffraction-limited spot of 200 nm diameter. Because the phases diffuse slowly on SLBs, a location on the SLB could be assessed by diffraction-limited fluorescence imaging to be *L_d_*, *L_o_*, or a homogeneous *L* phase before performing FCS. FCS revealed the diffusion of DPPE-TR to be 2x faster for POPC than DiPhyPC SLBs; *D_FCS_* = 4.9 ± 0.2 μm^2^/s and 2.5 ± 0.8 μm^2^/s, respectively (Table S2). *D_FCS_* was faster for the *L_d_* vs. *L_o_* phases, although not significantly for the cholesterol-rich 1:1:2 SLB; *D_FCS_* was (1.6 ± 0.5)x faster for *L_d_* vs. *L_o_* phases in the 2:2:1 SLBs and (1.1 ± 0.4)x for the 1:1:2 SLBs. This result is consistent in demonstrating is a greater difference in diffusion between the *L_d_* and *L_o_* phases for a membrane that has less cholesterol and a longer tie-line.

### 3.6) Diffusion vs. temperature

Increasing the sample temperature increased the single-lipid diffusion in all tested conditions (Fig. 6). Change in the diffusion vs. temperature was fit by a free area model derived from the kinetic theory of gas, which assumes that diffusion occurs when lipids hop to a surrounding transient void or free area that is created by a thermal density fluctuation (Galla et al., 1979). Since lipid hopping is an activated process dominated by van der Waal’s interactions, activation energy (*E_A_*) represents the energy barrier to be overcome for hopping between initial and final states that limits the molecular-scale lipid translation (Filippov et al., 2003; Macedo and Litovitz, 1965; Vaz et al., 1985). Thus, the diffusion coefficient vs. temperature can be described by the Arrhenius equation,

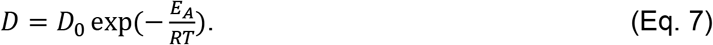

*E_A_* represents the activation energy, and *D_0_* represents the diffusion rate at very high temperatures. Fitting this exponential yielded temperature-independent values for *E_A_* and *D_0_* that are consistent with prior studies (Fig. 6 and Table S3).

**Figure 6:**
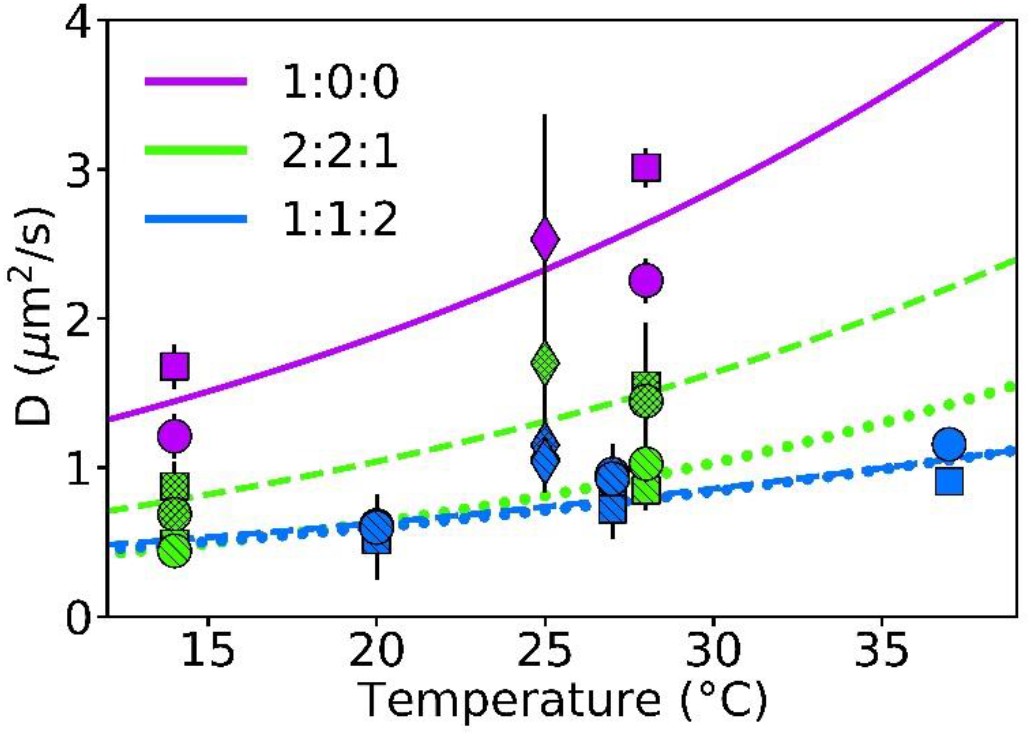
The diffusion of DPPE-TR was measured with varying membrane composition, lipid phase, temperature, and measurement technique. The color indicates the DiPhyPC:DPPC:cholesterol molar ratios. The marker shape indicates the measurement technique: square = *D_MSD_*; circle = *D_RD_*; diamond = *D_FCS_*. The marker hatching indicates the lipid phase: backslash = *L_o_*; crosshatching = *L_d_*; empty = *L or L_α_*. Fits of Eq. 7 are shown for each phase (Table S3). Error bars represent the standard error of repeated measurements with at least three repeats.

## 4) DISCUSSION

### 4.1) Phase separation on SLBs

Unlike domains on GUVs, all resolvable lipid phase in SLBs were immobile on the time scale of our experiments (<2 hr). The interaction between the glass substrate and the SLB was too strong for optically resolvable domains to move or coalescence, consistent with previous observations (Gunderson and Honerkamp-Smith, 2018). The immobile domains were convenient for correlating single-lipid diffusion with phase boundaries due to the minimal diffusion of the phases boundary during SPT data acquisition; however, single-lipid diffusion itself is also affected by and the membrane-substrate interaction. Lipids in GUVs diffuse 3x faster than lipids in SLBs (Beckers et al., 2020), and DPPE-TR only in the top leaflet was (1.2 ± 0.1)x faster than when in both leaflets (Woodward et al., 2018). If there was an equal contribution from DPPE-TR in both leaflets of the SLB, then the DPPE-TR in the top leaflet diffused 1.5x faster than DPPE-TR in the bottom leaflet.

Regardless of the initial domain distribution, we did not observe large domains (radius > 1 μm) form upon cooling well-mixed SLBs. The domains from well-mixed SLBs grew up to a size near the diffraction-limited imaging resolution. The domains were clearly present, but their sizes were difficult to determine due to both optical blurring and their non-circular shape. Substrate effects raise *T_m_* with a leaflet and lipid-type dependency such that not all lipids were well mixed after heating even when a uniform fluorescence emission was observed (Gunderson and Honerkamp-Smith, 2018). This effect may be amplified when Ca^2+^ is present and lipid pinning sites occur.

The slower temperature changes and increased duration at warm temperatures decreased the spatial correlation of domain locations upon thermal cycling, as shown previously (Stanich et al., 2013). However, the substrate-membrane interaction consistently caused all optically resolvable domains to be immobile and prevented their growth by domain merger. Additionally, the surface roughness of the substrate may have increased the domain nucleation rate (Goodchild et al., 2019), resulting in nanoscale domains present across the membrane instead of macroscale domains forming by gradual accretion. Phase-separated SLBs domains on glass or nanoscale roughened mica are orders of magnitude smaller than on mica; the membrane domain size was more correlated to substrate roughness rather than surface chemistry (Goodchild et al., 2019).

### 4.2) Causes of lipid aggregation

As described above, lipid aggregation may occur due to a variety of mechanisms. For example, rhodamine-labeled lipids form oligomers and grow into nanodomains due to the interactions of their extended aromatic moiety (Jan Akhunzada et al., 2019), which may be enhanced by membrane curvature (Woodward et al., 2018). The Texas Red-labeled lipids used here are similar in fluorophore structure with rhodamine-labeled lipids. Addtionally, the saturated acyl tailes of the DPPE-TR has been correlted with nanodomains within extended lipid phases (Sodt et al., 2014; Wu et al., 2016).

Light is capable of inducing phase separations via lipid peroxidation that is accelerated by fluorophores (Ayuyan and Cohen, 2006; Zhao et al., 2007). For example, some fluorophores such as Bodipy, DiO, DiI, Texas Red, and napthopyrene cause light-induced chemical changes to the lipids, including the oxidization of the unsaturated acyl tails (Zhao et al., 2007). Accordingly, DiPhyPC is frequently used for fluorescence-based diffusion studies because it provides a highly disordered acyl structure without carbon-carbon double bonds and resists light-induced chemical changes (Lindsey et al., 1979). DPPE-TR and light-induced aggregates are non-linearly dependent on the DPPE-TR concentration. Domain formation was 50x faster when 0.8 mol% rather than 0.15 mol% DPPE-TR was present (Zhao et al., 2007). The 15 mW of 561 nm wavelength light used here for 3 min of observation in the presence of 0.1 mol% DPPE-TR would be sufficient to cause light-induced alterations to the membrane phase behavior in the presence of unsaturated phospholipids. Our use of DiPhyPC was designed to minimize these light effects.

Lipid aggregates were previously seen more abundant in *L_o_* domains (Wu et al., 2016), yet no correlation between lipid phases and aggregation was observed here. In neither the 1:1:2 nor 2:2:1 SLBs was there a difference in the aggregation between the *L_d_* and *L_o_* phases. However, the 1:1:2 SLBs demonstrated greater aggregation than the 2:2:1 SLBs at all temperatures (Fig. 4). The parameters that determine aggregate formation are complex and depend on more than the lipid phase. For example, the greater abundance of aggregates in the DiPhyPC vs. POPC bilayers shown here are of unknown cause and worthy of further examination.

### 4.3) Diffusion of lipid aggregates

Single-lipid diffusion provides a detailed examination of the behavior of individual molecules with the possibility to resolve variations between diffusion modes, sample heterogeneity, and non-Brownian behaviors. Confined single-molecule trajectories were observed here coincident with lipid aggregates. When single-particle tracking was previously performed with long trajectories (*i.e*., over 100 steps per trajectory), the downward curvature of the MSD vs. Δ*t*, local variations in the diffusion rate, or transient spatial confinement was directly observed to reveal aggregation (Simson et al., 1995; Wu et al., 1977). Trajectories of fewer steps are amendable to detecting aggregates through analysis of the histogram of single-step lengths and comparison to Rayleigh distributions (Eq. 4). Aggregates displayed shorter step lengths than freely diffusing lipids, and the histogram of step lengths were fit best when multiple diffusers are assumed to be present (Spillane et al., 2014). Multiple-population fitting can yield the relative speed and abundance of each type of diffuser; however, single-step analysis requires each population to be >5% of the total and provide >10x difference in diffusion rates for reliable separation of the populations. Since this was not present in SLBs examined here, we alternatively relied on the varying localization rate for aggregates vs. the surrounding SLB to identify and exclude the aggregates from further analysis.

### 4.4) Resolution of Rayleigh distribution vs. MSD analyses

MSD and Rayleigh distribution analyses provided different precision in revealing the spatial resolution of diffusion differences. The MSD analysis typically provides greater accuracy in finding *D*, distributions in *D* from heterogenious populations of diffusers, and inherently correction for the localization uncertainty when not assuming MSD = 0 when *Δt* = 0 (Kabbani et al., 2020). Rayleigh distribution analysis, however, typically provide improved resolution in spatially varying *D*, assuming the single-particle trajectories are dense. For example, Rayleigh distribution analyses have been used to reveal the diffusion in 25 nm bins surrounding nanoscale membrane curvature (Kabbani et al., 2017; Kabbani and Kelly, 2017a; Woodward et al., 2018; Woodward and Kelly, 2020).

To demonstrate the tradeoff between spatial resolution and precision in determining *D*, let us consider a hypothetical diffuser measured by SPT with varying analysis methods. If a Brownian diffuser of *D* = 1 μm^2^/s was tracked for 1 sec, then the diffuser would be expected to traverse 2 μm, which would result in an averaging of spatial variations in diffusion across this distance for a typical single-molecule MSD analysis. However, spatial resolution from Rayleigh distribution analysis depends on the acquisition frame rate and the length of each step with an inherent averaging over this length. If the acquisition occurs at 100 Hz, then the step length and spatial resolution would be for Rayleigh distribution analysis would be 200 nm. This example demonstrates a 10x improvement in spatial resolution for Rayleigh distribution vs. MSD analysis. For short trajectories, such those used in this study, the MSD from each trajectory has large uncertainty; combining the trajectories from multiple molecules is necessary for accurate diffusion measurements for either MSD or Rayleigh distribution analysis, which further reduces the benefits of the MSD analysis.

The certainty with which *D* is determined can be approximated through statistical considerations. Both Rayleigh distribution and MSD analyses are Poisson noise dominated process such that the certainty in *D* is inversely proprotaional to the square root of the number of steps analyzed. Any gains in the area spatial resolution are matched by a proportional decrease in precision determining *D*. Unlike traditional MSD analyses, Rayleigh distribution fitting can be performed by averaging the step lengths over whatever area of the sample is warranted by the experimental details with this trade-off in mind.

### 4.5) Comparison between *D_MSD_, D_RD_*, and *D_FCS_*

*D_MSD_* and *D_RD_* were compared for all membrane composition, temperature, and phases (Fig. 6). *D_MSD_* and *D_RD_* varied by 18 ± 9%, which is typically within the experimental uncertainty for measuring the diffusion coefficient by any single method. *D_MSD_* and *D_RD_* displayed consistent trends in comparing lipid phases and varying temperatures. There were consistent differences between *D_MSD_* and *D_RD_* that depended on the rate of diffusion. Typically, when *D_MSD_* was > 1 μm^2^/s, *D_MSD_* was 1.2 ± 0.1 times larger than *D_RD_*, and otherwise *D_MSD_* was less than *D_RD_*. This systematic variation between *D_MSD_* and *D_RD_* remains unexplained but could be a coincidence in our analyses.

Confocal FCS consistently provides a resolution consistent with the difffration limit (*i.e*., 200 nm), but is highly limited in its detection of subpopulations. *D_FCS_* of DPPE-TR in POPC was 1.5 ± 0.5 times faster than *D_MSD_* and *D_RD_* (Tables S1 and S2). Commonly, FCS reports faster diffusion than SPT studies because *D_FCS_* does not incorporate immobile or highly confined diffusive subpopulations, whereas SPT typically does. Even after aggregate removal, SPT yield slower diffusion than FCS, but the differences were within the measured uncertainty of D.

### 4.6) Diffusion differences between lipid phases

When the tie-line between coexisting phases was longer, the difference between their diffusion was greater. The high cholesterol and short tie-line 1:1:2 SLBs displayed no significant difference in the diffusion rate between the *L_d_* and *L_o_* phases. However, the *L_d_* phase of 2:2:1 SLBs displayed diffusion twice as fast as the *L_o_* in 1:1:2 SLBs. The *L_o_* phases in the 1:1:2 and 2:2:1 SLBs did not display significantly different diffusion. Accordingly, increasing cholesterol content slowed the diffusion coefficient for the *L_d_* phase but not the *L_o_* phase. This is in contrast to ternary mixtures of dioleoylphosphatidylcholine, sphingomyelin, and cholesterol that display greater cholesterol content is correlated with slower diffusion in the *L_o_* phases and minimally changes in *L_d_* phase (Kahya et al., 2003; Scherfeld et al., 2003).

The 2:2:1 SLBs displayed faster DPPE-TR diffusion in *L_d_* vs. *L_o_* at ratios similar to seen previously (Dietrich et al., 2001). When the cholesterol content increased to 50% of the bilayer, (*i.e*., in 1:1:2 SLBs), the difference between *L_d_* and *L_o_* diffusion became indistinguishable for all tested temperatures. This is consistent with prior work that showed single population diffusion on GUVs when the increasing cholesterol content induced phase miscibility (Scherfeld et al., 2003). Increasing the cholesterol content for phase-separated membranes causes a shortening of the tie-line and increasing the similarity between the *L_o_* and *L_d_* phases until the membrane composition no longer supports coexisting phases. Interestingly, the aggregates diffusive properties are not significantly different between *L_d_* and *L_o_* of 2:21 SLBs (Woodward and Kelly, 2020).

### 4.7) Temperature affects diffusion

The diffusion of DPPE-TR was faster at higher temperatures for all compositions tested, similar to as shown previously (Bag et al., 2014; Filippov et al., 2004; Sengupta et al., 2008; Tamm, 1988). The changes in *D* with temperature was robust to SLB composition or phase. The data acquired here were not sufficient to significantly distinguish between the *D_0_* and *E_a_* values of each composition. However, it is interesting to note that the *E_a_* for the *L_o_* phase was (1.07 ± 0.03)x higher than the *L_d_* phase for both the 1:1:2 and 2:2:1 SLBs. Similarly, the *E_a_* values for the 1:1:2 SLBs were (1.5 ± 0.8)x higher than the 2:2:1 SLBs. These results make intuitive sense in that a greater barrier exists for lipids to exchange locations in a *L_o_* vs. *L_d_* phases or with higher cholesterol content.

## 5) CONCLUSIONS

This manuscript reports the effects of lipid composition, phase, temperature, and data analysis procedures on the single-lipid diffusion in model membranes. The importance of the tie-line length separating coexisting liquid phases was repeatedly demonstrated to be a key variable in predicting the differences between the *L_d_* and *L_o_* phases. The tie-lines shorten with increasing temperature and increasing cholesterol content as the lipid diffusion becomes more similar within the phases. Single-lipid diffusion was consistently faster at higher temperatures and was consistent with a free area model for diffusion in which higher cholesterol and ordered phases resulted in a greater activation energy for laterial lipid diffusion.

DiPhyPC-containing membranes displayed nanoscale fluorescent lipid aggregation in both *L_o_* and *L_d_* domains. These aggregations were culled via localization rate thresholding. The single-lipid diffusion was analyzed with and without the aggregates culled. Increasing cholesterol and decreasing temperature both correlated with greater aggregation. However, there was no significant correlation between lipid phase and aggregation. Single-particle tracking data was analyzed via Rayleigh distributions and MSD analyses. Rayleigh distribution analyses have yield improved spatial resolution in heterogeneous samples. The protocols reported here demonstrate consistency between complementary methods of measuring lipid diffusion while detailing the advantages of each method.

## 6) ACKNOWLEDGMENTS

The authors thank Aurelia R. Honerkamp-Smith for valuable discussions. Financial support was provided by Thomas C. Rumble University graduate fellowship, Wayne State University summer dissertation award, and Richard J. Barber. This material is based upon work supported by the National Science Foundation under grant no. DMR-1652316.

## 7) ABBREVIATIONS

DiPhyPC: diphytanoylphosphatidylcholine
DPPC: dipalmitoylphosphatidylcholine
DPPE-TR: dipalmitoylphosphoethanolamine-Texas Red
FCS: fluorescent correlation spectroscopy
GUV: giant unilamellar vesicle
L_α_: liquid phase of a one component membrane
L_d_: liquid disordered phase
L_o_: liquid ordered phase
L: liquid phase
MSD: mean squared displacement
*N*: number of localizations per aggregate
POPC: palmitoyloleoylphosphatidylcholine
SLB: supported lipid bilayer
SPT: single-particle tracking

## SUPPLEMENTAL MATERIAL

**Table S1:** SPT diffusion results via MSD vs. Δt analysis after aggregate removal.

| Composition | Temperature (°C) | Phase | $D_{RD} (\mu\text{m}^2/\text{s})$ | $D_{MSD} (\mu\text{m}^2/\text{s})$ |
| --- | --- | --- | --- | --- |
| POPC | 25 | $L_\alpha$ | $3.1 \pm 0.4$ | $3.2 \pm 1.2$ |
| 1:0:0 | 14 | $L_\alpha$ | $1.2 \pm 0.2$ | $1.7 \pm 0.2$ |
| 1:0:0 | 28 | $L_\alpha$ | $2.3 \pm 0.2$ | $3.0 \pm 0.1$ |
| 2:2:1 | 14 | $L_d$ | $0.7 \pm 0.1$ | $0.9 \pm 0.2$ |
| 2:2:1 | 14 | $L_o$ | $0.4 \pm 0.1$ | $0.5 \pm 0.1$ |
| 2:2:1 | 28 | $L_d$ | $1.5 \pm 0.2$ | $1.6 \pm 0.4$ |
| 2:2:1 | 28 | $L_o$ | $1.0 \pm 0.1$ | $0.9 \pm 0.1$ |
| 1:1:2 | 20 | $L_d$ | $0.6 \pm 0.2$ | $0.5 \pm 0.3$ |
| 1:1:2 | 20 | $L_o$ | $0.6 \pm 0.2$ | $0.5 \pm 0.2$ |
| 1:1:2 | 27 | $L_d$ | $1.0 \pm 0.2$ | $0.8 \pm 0.3$ |
| 1:1:2 | 27 | $L_o$ | $0.9 \pm 0.1$ | $0.7 \pm 0.1$ |
| 1:1:2 | 37 | $L$ | $1.2 \pm 0.1$ | $0.9 \pm 0.1$ |

**Table S2:** FCS diffusion results.

| Composition | Temperature (°C) | Phase | $D_{FCS} (\mu\text{m}^2/\text{s})$ |
| --- | --- | --- | --- |
| POPC | 25 | $L_\alpha$ | $4.9 \pm 0.2$ |
| 1:0:0 | 25 | $L_\alpha$ | $2.5 \pm 0.8$ |
| 2:2:1 | 25 | $L_d$ | $1.7 \pm 0.5$ |
| 2:2:1 | 25 | $L_o$ | $1.1 \pm 0.1$ |
| 1:1:2 | 25 | $L_d$ | $1.2 \pm 0.3$ |
| 1:1:2 | 25 | $L_o$ | $1.0 \pm 0.2$ |

**Table S3:** *D_0_* and *E_A_* fit results. Uncertainties are extracted from the covariance matrix resulting from the least squares fitting of Eq. 7, as shown in Fig. 6.

| Composition | Phase | $E_A$ (kJ/mol) | $D_0$ (mm<sup>2</sup>/s) |
| --- | --- | --- | --- |
| 1:0:0 | $L_\alpha$ | $31 \pm 13$ | $0.59 \pm 3$ |
| 2:2:1 | $L_d$ | $23 \pm 7$ | $0.008 \pm 0.02$ |
| 2:2:1 | $L_o$ | $24 \pm 7$ | $0.016 \pm 0.04$ |
| 1:1:2 | $L_d$ | $33 \pm 6$ | $0.94 \pm 2$ |
| 1:1:2 | $L_o$ | $36 \pm 8$ | $1.5 \pm 5$ |

## Notes

https://www.cvkelly.com

### Competing Interest Statement

The authors have declared no competing interest.

## REFERENCES

Almeida, P.F.F., Vaz, W.L.C., Thompson, T.E., 1992. Lateral diffusion and percolation in two-phase, two-component lipid bilayers. Topology of the solid-phase domains in-plane and across the lipid bilayer. Biochemistry 31, 7198–7210. https://doi.org/10.1021/bi00146a024

Ayuyan, A.G., Cohen, F.S., 2006. Lipid Peroxides Promote Large Rafts: Effects of Excitation of Probes in Fluorescence Microscopy and Electrochemical Reactions during Vesicle Formation. Biophys J 91, 2172–2183. https://doi.org/10.1529/biophysj.106.087387

Bag, N., Yap, D.H.X., Wohland, T., 2014. Temperature dependence of diffusion in model and live cell membranes characterized by imaging fluorescence correlation spectroscopy. Biochim. Biophys. Acta 1838, 802–813. https://doi.org/10.1016/j.bbamem.2013.10.009

Baumgart, T., Hess, S.T., Webb, W.W., 2003. Imaging coexisting fluid domains in biomembrane models coupling curvature and line tension. Nature 425, 821–824. https://doi.org/10.1038/nature02013

Beckers, D., Urbancic, D., Sezgin, E., 2020. Impact of Nanoscale Hindrances on the Relationship between Lipid Packing and Diffusion in Model Membranes. J. Phys. Chem. B. https://doi.org/10.1021/acs.jpcb.0c00445

Berglund, A.J., 2010. Statistics of camera-based single-particle tracking. Phys. Rev. E 82, 011917. https://doi.org/10.1103/PhysRevE.82.011917

Bleecker, J.V., Cox, P.A., Foster, R.N., Litz, J.P., Blosser, M.C., Castner, D.G., Keller, S.L., 2016. Thickness Mismatch of Coexisting Liquid Phases in Non-Canonical Lipid Bilayers. J Phys Chem B 120, 2761–2770. https://doi.org/10.1021/acs.jpcb.5b10165

Cheney, P.P., Weisgerber, A.W., Feuerbach, A.M., Knowles, M.K., 2017. Single Lipid Molecule Dynamics on Supported Lipid Bilayers with Membrane Curvature. Membranes (Basel) 7. https://doi.org/10.3390/membranes7010015

Chiantia, S., Kahya, N., Ries, J., Schwille, P., 2006a. Effects of Ceramide on Liquid-Ordered Domains Investigated by Simultaneous AFM and FCS. Biophys J 90, 4500–4508. https://doi.org/10.1529/biophysj.106.081026

Chiantia, S., Ries, J., Kahya, N., Schwille, P., 2006b. Combined AFM and two-focus SFCS study of raft-exhibiting model membranes. Chemphyschem 7, 2409–2418. https://doi.org/10.1002/cphc.200600464

Coker, H.L.E., Cheetham, M.R., Kattnig, D.R., Wang, Y.J., Garcia-Manyes, S., Wallace, M.I., 2019. Controlling Anomalous Diffusion in Lipid Membranes. Biophys J 116, 1085–1094. https://doi.org/10.1016/j.bpj.2018.12.024

Dietrich, C., Bagatolli, L.A., Volovyk, Z.N., Thompson, N.L., Levi, M., Jacobson, K., Gratton, E., 2001. Lipid rafts reconstituted in model membranes. Biophys J 80, 1417–1428.

Dimova, R., 2014. Recent developments in the field of bending rigidity measurements on membranes. Advances in Colloid and Interface Science, Special issue in honour of Wolfgang Helfrich 208, 225–234. https://doi.org/10.1016/j.cis.2014.03.003

Fessler, M.B., Parks, J.S., 2011. Intracellular lipid flux and membrane microdomains as organizing principles in inflammatory cell signaling. J. Immunol. 187, 1529–1535. https://doi.org/10.4049/jimmunol.1100253

Filippov, A., Orädd, G., Lindblom, G., 2004. Lipid Lateral Diffusion in Ordered and Disordered Phases in Raft Mixtures. Biophysical Journal 86, 891–896. https://doi.org/10.1016/S0006-3495(04)74164-8

Filippov, A., Orädd, G., Lindblom, G., 2003. The Effect of Cholesterol on the Lateral Diffusion of Phospholipids in Oriented Bilayers. Biophys J 84, 3079–3086.

Galla, H.-J., Hartmann, W., Theilen, U., Sackmann, E., 1979. On two-dimensional passive random walk in lipid bilayers and fluid pathways in biomembranes. J. Membrain Biol. 48, 215–236. https://doi.org/10.1007/BF01872892

Goodchild, J.A., Walsh, D.L., Connell, S.D.A., 2019. Nanoscale Substrate Roughness Hinders Domain Formation in Supported Lipid Bilayers. Langmuir. https://doi.org/10.1021/acs.langmuir.9b01990

Gracià, R.S., Bezlyepkina, N., Knorr, R.L., Lipowsky, R., Dimova, R., 2010. Effect of cholesterol on the rigidity of saturated and unsaturated membranes: fluctuation and electrodeformation analysis of giant vesicles. Soft Matter 6, 1472–1482. https://doi.org/10.1039/B920629A

Gunderson, R.S., Honerkamp-Smith, A.R., 2018. Liquid-liquid phase transition temperatures increase when lipid bilayers are supported on glass. Biochimica et Biophysica Acta (BBA) - Biomembranes, Emergence of Complex Behavior in Biomembranes 1860, 1965–1971. https://doi.org/10.1016/j.bbamem.2018.05.001

Guo, S.-M., He, J., Monnier, N., Sun, G., Wohland, T., Bathe, M., 2012. Bayesian approach to the analysis of fluorescence correlation spectroscopy data II: application to simulated and in vitro data. Anal. Chem. 84, 3880–3888. https://doi.org/10.1021/ac2034375

Harwardt, M.-L.I.E., Dietz, M.S., Heilemann, M., Wohland, T., 2018. SPT and Imaging FCS Provide Complementary Information on the Dynamics of Plasma Membrane Molecules. Biophys J 114, 2432–2443. https://doi.org/10.1016/j.bpj.2018.03.013

Hsieh, C.-L., Spindler, S., Ehrig, J., Sandoghdar, V., 2014. Tracking single particles on supported lipid membranes: multimobility diffusion and nanoscopic confinement. J Phys Chem B 118, 1545–1554. https://doi.org/10.1021/jp412203t

Hurley, J.H., Boura, E., Carlson, L.-A., Rózycki, B., 2010. Membrane Budding. Cell 143, 875–887. https://doi.org/10.1016/j.cell.2010.11.030

Jacobson, K., Hou, Y., Derzko, Z., Wojcieszyn, J., Organisciak, D., 1981. Lipid lateral diffusion in the surface membrane of cells and in multibilayers formed from plasma membrane lipids. Biochemistry 20, 5268–5275. https://doi.org/10.1021/bi00521a027

Jan Akhunzada, M., D’Autilia, F., Chandramouli, B., Bhattacharjee, N., Catte, A., Di Rienzo, R., Cardarelli, F., Brancato, G., 2019. Interplay between lipid lateral diffusion, dye concentration and membrane permeability unveiled by a combined spectroscopic and computational study of a model lipid bilayer. Sci Rep 9, 1508. https://doi.org/10.1038/s41598-018-37814-x

Jaqaman, K., Loerke, D., Mettlen, M., Kuwata, H., Grinstein, S., Schmid, S.L., Danuser, G., 2008. Robust single-particle tracking in live-cell time-lapse sequences. Nat Meth 5, 695–702. https://doi.org/10.1038/nmeth.1237

Kabbani, A.M., Kelly, C.V., 2017a. The Detection of Nanoscale Membrane Bending with Polarized Localization Microscopy. Biophysical Journal 113, 1782–1794. https://doi.org/10.1016/j.bpj.2017.07.034

Kabbani, A.M., Kelly, C.V., 2017b. Nanoscale Membrane Budding Induced by CTxB and Detected via Polarized Localization Microscopy. Biophysical Journal 113, 1795–1806. https://doi.org/10.1016/j.bpj.2017.08.031

Kabbani, A.M., Raghunathan, K., Lencer, W.I., Kenworthy, A.K., Kelly, C.V., 2020. Structured clustering of the glycosphingolipid GM1 is required for membrane curvature induced by cholera toxin. bioRxiv 2020.01.22.915249. https://doi.org/10.1101/2020.01.22.915249

Kabbani, A.M., Woodward, X., Kelly, C.V., 2017. Revealing the Effects of Nanoscale Membrane Curvature on Lipid Mobility. Membranes 7, 60. https://doi.org/10.3390/membranes7040060

Kahya, N., Scherfeld, D., Bacia, K., Poolman, B., Schwille, P., 2003. Probing lipid mobility of raft-exhibiting model membranes by fluorescence correlation spectroscopy. J. Biol. Chem. 278, 28109–28115. https://doi.org/10.1074/jbc.M302969200

Kiessling, V., Yang, S.-T., Tamm, L.K., 2015. Supported lipid bilayers as models for studying membrane domains. Curr Top Membr 75, 1–23. https://doi.org/10.1016/bs.ctm.2015.03.001

Knight, J.D., Lerner, M.G., Marcano-Velázquez, J.G., Pastor, R.W., Falke, J.J., 2010. Single Molecule Diffusion of Membrane-Bound Proteins: Window into Lipid Contacts and Bilayer Dynamics. Biophysical Journal 99, 2879–2887. https://doi.org/10.1016/j.bpj.2010.08.046

Kollmitzer, B., Heftberger, P., Podgornik, R., Nagle, J.F., Pabst, G., 2015. Bending Rigidities and Interdomain Forces in Membranes with Coexisting Lipid Domains. Biophys. J. 108, 2833–2842. https://doi.org/10.1016/j.bpj.2015.05.003

Lagerholm, B.C., Andrade, D.M., Clausen, M.P., Eggeling, C., 2017. Convergence of lateral dynamic measurements in the plasma membrane of live cells from single particle tracking and STED-FCS. J Phys D Appl Phys 50, 063001. https://doi.org/10.1088/1361-6463/aa519e

Lin, W.-C., Blanchette, C.D., Longo, M.L., 2007. Fluid-Phase Chain Unsaturation Controlling Domain Microstructure and Phase in Ternary Lipid Bilayers Containing GalCer and Cholesterol. Biophys J 92, 2831–2841. https://doi.org/10.1529/biophysj.106.095422

Lindblom, G., Orädd, G., Filippov, A., 2006. Lipid lateral diffusion in bilayers with phosphatidylcholine, sphingomyelin and cholesterol: An NMR study of dynamics and lateral phase separation. Chemistry and Physics of Lipids, Protein:Lipid Interactions 141, 179–184. https://doi.org/10.1016/j.chemphyslip.2006.02.011

Lindsey, H., Petersen, N.O., Chan, S.I., 1979. Physicochemical characterization of 1,2-diphytanoyl-sn-glycero-3-phosphocholine in model membrane systems. Biochim. Biophys. Acta 555, 147–167. https://doi.org/10.1016/0005-2736(79)90079-8

Macedo, P.B., Litovitz, T.A., 1965. On the Relative Roles of Free Volume and Activation Energy in the Viscosity of Liquids. J. Chem. Phys. 42, 245–256. https://doi.org/10.1063/1.1695683

McConnell, H.M., Radhakrishnan, A., 2003. Condensed complexes of cholesterol and phospholipids. Biochim. Biophys. Acta 1610, 159–173. https://doi.org/10.1016/s0005-2736(03)00015-4

Ovesný, M., Křížek, P., Borkovec, J., Svindrych, Z., Hagen, G.M., 2014. ThunderSTORM: a comprehensive ImageJ plug-in for PALM and STORM data analysis and super-resolution imaging. Bioinformatics 30, 2389–2390. https://doi.org/10.1093/bioinformatics/btu202

Pike, L.J., 2006. Rafts defined: a report on the Keystone Symposium on Lipid Rafts and Cell Function. J. Lipid Res. 47, 1597–1598. https://doi.org/10.1194/jlr.E600002-JLR200

Pralle, A., Keller, P., Florin, E.-L., Simons, K., Hörber, J.K.H., 2000. Sphingolipid–Cholesterol Rafts Diffuse as Small Entities in the Plasma Membrane of Mammalian Cells. J Cell Biol 148, 997–1008.

Qian, H., Sheetz, M.P., Elson, E.L., 1991. Single particle tracking. Analysis of diffusion and flow in two-dimensional systems. Biophys J 60, 910–921.

Radhakrishnan, A., McConnell, H., 2005. Condensed complexes in vesicles containing cholesterol and phospholipids. Proc Natl Acad Sci U S A 102, 12662–12666. https://doi.org/10.1073/pnas.0506043102

Saxton, M.J., 1997. Single-particle tracking: the distribution of diffusion coefficients. Biophys J 72, 1744–1753.

Scherer, K.M., Spille, J.-H., Sahl, H.-G., Grein, F., Kubitscheck, U., 2015. The Lantibiotic Nisin Induces Lipid II Aggregation, Causing Membrane Instability and Vesicle Budding. Biophysical Journal 108, 1114–1124. https://doi.org/10.1016/j.bpj.2015.01.020

Scherfeld, D., Kahya, N., Schwille, P., 2003. Lipid Dynamics and Domain Formation in Model Membranes Composed of Ternary Mixtures of Unsaturated and Saturated Phosphatidylcholines and Cholesterol. Biophysical Journal 85, 3758–3768. https://doi.org/10.1016/S0006-3495(03)74791-2

Schindelin, J., Arganda-Carreras, I., Frise, E., Kaynig, V., Longair, M., Pietzsch, T., Preibisch, S., Rueden, C., Saalfeld, S., Schmid, B., Tinevez, J.-Y., White, D.J., Hartenstein, V., Eliceiri, K., Tomancak, P., Cardona, A., 2012. Fiji: an open-source platform for biological-image analysis. Nat Meth 9, 676–682. https://doi.org/10.1038/nmeth.2019

Sengupta, P., Hammond, A., Holowka, D., Baird, B., 2008. Structural Determinants for Partitioning of Lipids and Proteins Between Coexisting Fluid Phases in Giant Plasma Membrane Vesicles. Biochim Biophys Acta 1778, 20–32. https://doi.org/10.1016/j.bbamem.2007.08.028

Shelby, S.A., Holowka, D., Baird, B., Veatch, S.L., 2013. Distinct Stages of Stimulated FcεRI Receptor Clustering and Immobilization Are Identified through Superresolution Imaging. Biophys J 105, 2343–2354. https://doi.org/10.1016/j.bpj.2013.09.049

Simons, K., Ikonen, E., 1997. Functional rafts in cell membranes. Nature 387, 569–572. https://doi.org/10.1038/42408

Simons, K., Toomre, D., 2000. Lipid rafts and signal transduction. Nat. Rev. Mol. Cell Biol. 1, 31–39. https://doi.org/10.1038/35036052

Simson, R., Sheets, E.D., Jacobson, K., 1995. Detection of temporary lateral confinement of membrane proteins using single-particle tracking analysis. Biophys. J. 69, 989–993. https://doi.org/10.1016/S0006-3495(95)79972-6

Sodt, A.J., Sandar, M.L., Gawrisch, K., Pastor, R.W., Lyman, E., 2014. The Molecular Structure of the Liquid-Ordered Phase of Lipid Bilayers. J. Am. Chem. Soc. 136, 725–732. https://doi.org/10.1021/ja4105667

Spillane, K.M., Ortega-Arroyo, J., de Wit, G., Eggeling, C., Ewers, H., Wallace, M.I., Kukura, P., 2014. High-Speed Single-Particle Tracking of GM1 in Model Membranes Reveals Anomalous Diffusion due to Interleaflet Coupling and Molecular Pinning. Nano Lett. 14, 5390–5397. https://doi.org/10.1021/nl502536u

Stanich, C.A., Honerkamp-Smith, A.R., Putzel, G.G., Warth, C.S., Lamprecht, A.K., Mandal, P., Mann, E., Hua, T.-A.D., Keller, S.L., 2013. Coarsening Dynamics of Domains in Lipid Membranes. Biophysical Journal 105, 444–454. https://doi.org/10.1016/j.bpj.2013.06.013

Štefl, M., Šachl, R., Humpolíčková, J., Cebecauer, M., Macháň, R., Kolářová, M., Johansson, L.B.-Å., Hof, M., 2012. Dynamics and Size of Cross-Linking-Induced Lipid Nanodomains in Model Membranes. Biophys J 102, 2104–2113. https://doi.org/10.1016/j.bpj.2012.03.054

Sun, H., Chen, L., Gao, L., Fang, W., 2015. Nanodomain Formation of Ganglioside GM1 in Lipid Membrane: Effects of Cholera Toxin-Mediated Cross-Linking. Langmuir 31, 9105–9114. https://doi.org/10.1021/acs.langmuir.5b01866

Tamm, L.K., 1988. Lateral diffusion and fluorescence microscope studies on a monoclonal antibody specifically bound to supported phospholipid bilayers. Biochemistry 27, 1450–1457. https://doi.org/10.1021/bi00405a009

Uppamoochikkal, P., Tristram-Nagle, S., Nagle, J.F., 2010. Orientation of tie-lines in the phase diagram of DOPC/DPPC/cholesterol model biomembranes. Langmuir 26, 17363–17368. https://doi.org/10.1021/la103024f

Vaz, W.L.C., Clegg, R.M., Hallmann, D., 1985. Translational diffusion of lipids in liquid crystalline phase phosphatidylcholine multibilayers. A comparison of experiment with theory. Biochemistry 24, 781–786. https://doi.org/10.1021/bi00324a037

Veatch, S.L., 2007. Electro-formation and fluorescence microscopy of giant vesicles with coexisting liquid phases. Methods Mol. Biol. 398, 59–72. https://doi.org/10.1007/978-1-59745-513-8_6

Veatch, S.L., Gawrisch, K., Keller, S.L., 2006. Closed-Loop Miscibility Gap and Quantitative Tie-Lines in Ternary Membranes Containing Diphytanoyl PC. Biophys J 90, 4428–4436. https://doi.org/10.1529/biophysj.105.080283

Veatch, S.L., Keller, S.L., 2005. Seeing spots: Complex phase behavior in simple membranes. Biochimica et Biophysica Acta (BBA) - Molecular Cell Research, Lipid Rafts: From Model Membranes to Cells 1746, 172–185. https://doi.org/10.1016/j.bbamcr.2005.06.010

Veatch, S.L., Keller, S.L., 2002. Organization in lipid membranes containing cholesterol. Phys. Rev. Lett. 89, 268101. https://doi.org/10.1103/PhysRevLett.89.268101

Veatch, S.L., Machta, B.B., Shelby, S.A., Chiang, E.N., Holowka, D.A., Baird, B.A., 2012. Correlation Functions Quantify Super-Resolution Images and Estimate Apparent Clustering Due to Over-Counting. PLOS ONE 7, e31457. https://doi.org/10.1371/journal.pone.0031457

Wawrezinieck, L., Rigneault, H., Marguet, D., Lenne, P.-F., 2005. Fluorescence Correlation Spectroscopy Diffusion Laws to Probe the Submicron Cell Membrane Organization. Biophys J 89, 4029–4042. https://doi.org/10.1529/biophysj.105.067959

Woodward, X., Kelly, C.V., 2020. Coexisting lipid phases alter lipid dynamics and sort on nanoscale membrane curvature. Pending submission to the BioRxiv and peer-review.

Woodward, X., Stimpson, E.E., Kelly, C.V., 2018. Single-lipid tracking on nanoscale membrane buds: The effects of curvature on lipid diffusion and sorting. Biochimica et Biophysica Acta (BBA) - Biomembranes, Emergence of Complex Behavior in Biomembranes 1860, 2064–2075. https://doi.org/10.1016/j.bbamem.2018.05.009

Wu, E., Jacobson, K., Papahadjopoulos, D., 1977. Lateral diffusion in phospholipid multibilayers measured by fluorescence recovery after photobleaching. Biochemistry 16, 3936–3941. https://doi.org/10.1021/bi00636a034

Wu, H.-M., Lin, Y.-H., Yen, T.-C., Hsieh, C.-L., 2016. Nanoscopic substructures of raft-mimetic liquid-ordered membrane domains revealed by high-speed single-particle tracking. Scientific Reports 6, 20542. https://doi.org/10.1038/srep20542

Yuan, C., Furlong, J., Burgos, P., Johnston, L.J., 2002. The size of lipid rafts: an atomic force microscopy study of ganglioside GM1 domains in sphingomyelin/DOPC/cholesterol membranes. Biophys J 82, 2526–2535.

Yuan, C., Johnston, L.J., 2001. Atomic force microscopy studies of ganglioside GM1 domains in phosphatidylcholine and phosphatidylcholine/cholesterol bilayers. Biophys J 81, 1059–1069.

Zhao, J., Wu, J., Shao, H., Kong, F., Jain, N., Hunt, G., Feigenson, G., 2007. Phase studies of model biomembranes: Macroscopic coexistence of Lα + Lβ, with light-induced coexistence of Lα + Lo Phases. Biochim Biophys Acta 1768, 2777–2786. https://doi.org/10.1016/j.bbamem.2007.07.009

